# Comparative Analysis of Genetically-Modified Crops: Conditional Equivalence Criteria

**DOI:** 10.1101/2021.02.19.431950

**Authors:** Changjian Jiang, Chen Meng, Adam W. Schapaugh, Huizhe Jin

## Abstract

The comparative assessment of genetically-modified (GM) crops relies on the principle of substantial equivalence, which states that such products should be compared to conventional counterparts that have an established history of safe use. In an effort to operationalize this principle, the GMO Panel of the European Food Safety Authority proposed an equivalence test that directly compares a GM test variety with a set of unrelated, conventionally-bred reference varieties with part of the difference as the known background of the test (the same as the given control). The criterion of the EFSA test, however, is defined solely by genotypic differences between the non-traited control and reference varieties (i.e. the background effect) while assuming the so-called GM trait effect as zero. As the outcome of an EFSA equivalence test is determined primarily by the similarity, or lack thereof, of the control and references, a conditional equivalence criterion is proposed in this investigation that focuses on “unintended” effects of a GM trait which is irrespective of the (random) genotypic value of a given control. The new criterion also includes a mean-scaled standard similar to the 80-125% rule for bioequivalence assessment practiced in the pharmaceutical industry as an alternative when the reference variation is zero or close to zero. In addition, optional criteria are proposed with a step-wise procedure to control the rate of false negatives (non-equivalence by chance) providing a comprehensive assessment under multiple comparisons. An application to maize grain composition data demonstrates that the conditional equivalence criterion provides effect-specific and more robust assessment of equivalence than the EFSA criterion did, especially for GM traits showing negligible or no unintended effects which are likely true for most traits in the current market.

## Introduction

The comparative assessment of foods derived from genetically-modified (GM) crops relies on the principle of substantial equivalence, which states that such products should be compared to conventional counterparts that have an established history of safe use but are not required to have zero difference from a near-isogenic control line absent of a GM trait in terms of “natural variations” [1–4]. In an effort to operationalize this principle, the GMO Panel of the European Food Safety Authority (EFSA 2010) proposed an equivalence criterion (thereafter called EFSA equivalence criterion or limits) that compares a GM test variety with a set of conventionally-bred references with part of the difference as the known genotypic background of the test (the same as the near-isogenic control line) [5,6]. Similar criteria have also appeared in the literature [7,8]. Nevertheless, Codex states that “in achieving the objective of conferring a specific target trait (intended effect) to a plant…, additional traits… could be lost or modified (unintended effects)” and “the safety assessment of foods derived from recombinant-DNA plants involves methods to identify and detect such unintended effects and procedures to evaluate their biological relevance and potential impact on food safety” [2]. As described above EFSA’s method requests an assessment of differences of the test from a set of references regardless of differences from the control. These differences would contain a trait effect if presents and, however, are certainly driven by genotypic values of the control resulting from conventional plant breeding (as described by Jiang et al. [9]). Thus, the result of EFSA equivalence testing in practice often is unrelated to the trait effect, which should be the sole focus of the comparative assessment, creating a series of discussions in the recent literature [7–15].

As the outcome of an EFSA equivalence test is determined primarily by the similarity, or lack thereof, of the control and references (Fig 1), a conditional equivalence criterion is proposed in this investigation that focuses on “unintended effects” of a GM trait irrespective of the (random) genotypic values of a given control.

**Fig 1.**
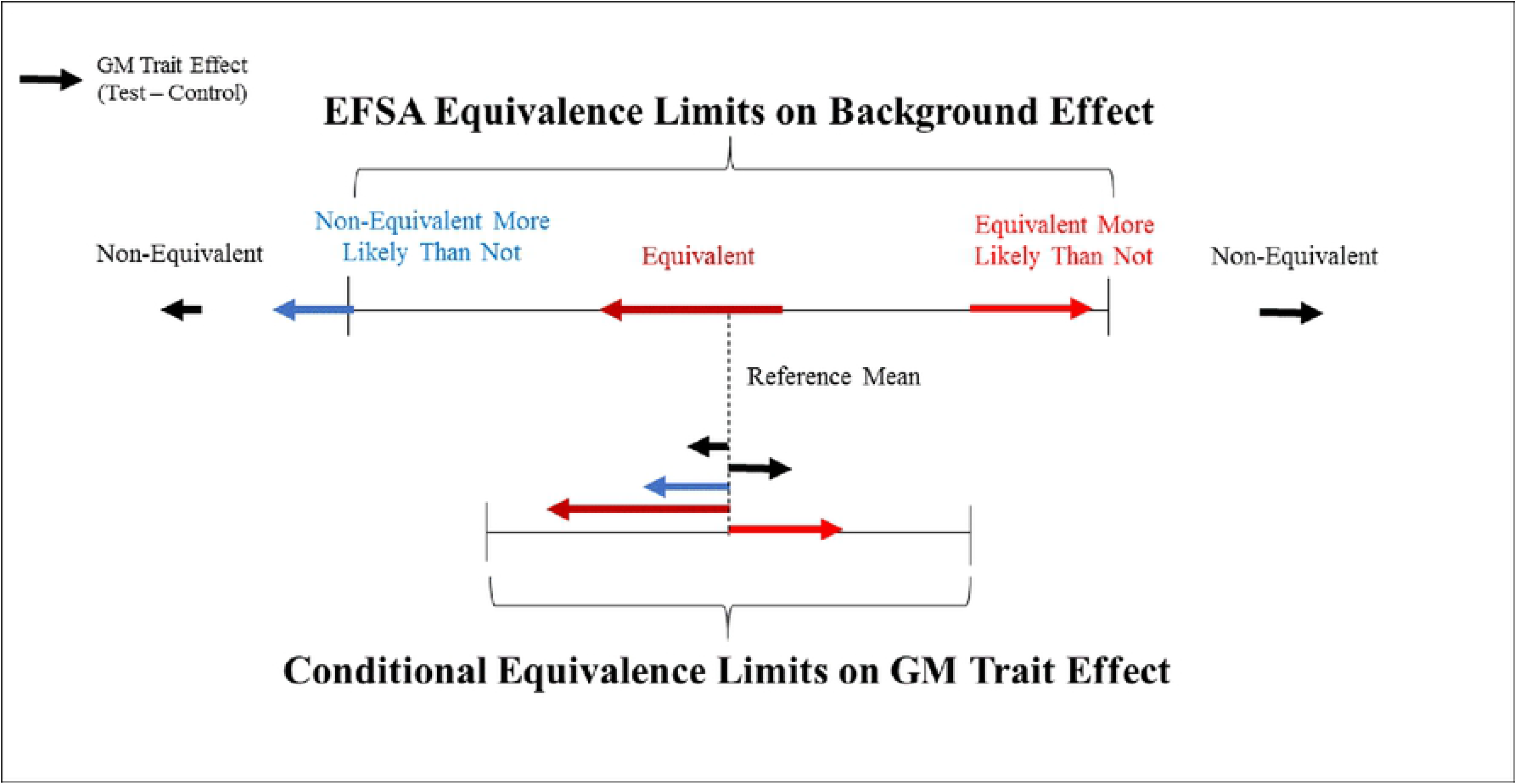
Graphical illustration of EFSA and conditional equivalence criteria. EFSA equivalence criterion is defined primarily by a (random) control background effect and has no specification of a GM trait effect, and a conditional equivalence criterion is defined solely for a GM trait effect with a given control.

To our knowledge, no equivalence criterion of a GM trait effect (with a given genotypic background) has been developed for the comparative assessment of GM crops and derived food/feed. All criteria in the literature are defined on the basis of the background effect assuming the absence of a GM trait effect [6,7,9] which have shown at least three limitations in practice: first is the sensitivity of the equivalence conclusion to a given control (i.e., the same GM product can generate different, completely contradictory, conclusions, based solely on the selected control) (Fig 1); second is the incomplete coverage of a one-size-fits-all criterion for a wide range of endpoints, with dramatic differences in their means and variations as explained in the next section; and third is the lack of a strategy for a control of false negatives (non-equivalence, in this case) in a comprehensive assessment under multiple comparisons as the criterion defined by a fixed percentile without an adjustment for the number of comparisons.

Here, a conditional equivalence criterion is derived on the basis of an expected mean squared difference of a GM crop from the references using a mixed model approach assuming the random background variation, similar to that used by the Food and Drug Administration (FDA) for individual bioequivalence in the presence of a random, individual-specific effect [16]. When the reference variation for the background effect is too low to provide a valid (variation-scaled) criterion, a mean-scaled criterion, similar to the 80-125% rule for the bioequivalence assessment in the pharmaceutical industry, is recommended as an alternative. Due to the alleviated false negative rate (much higher than the target level of 5% as the EFSA criterion defined by a 95% confidence interval), a data-driven procedure is proposed for selecting criteria with optional criteria to statistically control these errors. An application to a maize grain composition example (used by EFSA) demonstrates that the proposed conditional equivalence criteria provides substantial improvement for a true similarity measurement of GM crop over three limitations of EFSA criterion and others.

The organization of the manuscript starts with basic assumptions of the principle of substantial equivalence, followed by the derivation of a set of conditional equivalence criteria, and then a data-driven procedure for selecting criteria across various endpoints in practice, and finally an application to a maize grain composition example. The discussion includes additional thoughts on each of these new criteria, and highlights areas of further research.

## The definition of substantial equivalence

### Assumptions and the current practice

Assume a Test-Control-Reference (TCR) trial where the test is the GM variety, the control is an isogenic non-GM comparator with the genotypic background similar to the test (as monitored by the molecular breeding technique [17,18]), and the references are comparators with different genetic backgrounds. Let (*μ*_*T*_, *μ*_*C*_, *μ*_*R*_) denote parameters of genotypic group means, 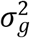 the genetic variation among references, and (Δ_*TC*_, Δ_*CR*_, Δ_*TR*_) the mean differences among three groups. For parameters of interest, Δ_*TC*_ represents a GM trait effect with a given control, while Δ_*CR*_ is an effect of the genotypic background shared by the test and the control from the traditional plant breeding. A simple difference of a GM test from the reference mean Δ_*TR*_ (= Δ_*TC*_ + Δ_*CR*_) as expected consists of both GM trait effect Δ_*TC*_ and background effect Δ_*CR*_.

The following are the underlying assumptions of the principle of substantial equivalence and the current global regulatory approval procedure of a GM crop.

a. GM crop safety assessment is solely interested in the effect of a GM trait regardless of the genotypic background; “It focuses on assessing the safety of any identified differences so that the safety of the new product can be considered relative to its conventional counterpart” [2]. Though the regulatory evaluation of a GM crop is on a given control background, upon approval, the GM trait could be integrated into any conventional reference (in the current market or from a breeding program) during the commercial application, and the background effect Δ_*CR*_ is expected to vary from endpoint to endpoint for a given control or for the same endpoint across different controls [17,18,19]. An equivalence of a GM crop should focus solely on a GM trait effect Δ_*TC*_ (= Δ_*TR*_ ― Δ_*CR*_) regardless of the genotypic background effect Δ_*CR*_ of a given control (Fig 1).
b. Substantial equivalence of a GM crop in statistics is a similarity measure to a distribution of conventional references with a history-of-safe-use; “Any observed differences should be assessed in the context of the range of natural variations” [2] demonstrated by conventional references with no requirement of a trait effect Δ_*TC*_ = 0. In spite of a given control was applied in a TCR trial, the equivalence of a GM crop in statistics is a similarity or distance measurement of the mean difference Δ_*TC*_ between two probability distributions, one for GM crop with various genotypic backgrounds and one for conventional references, in the scale of the reference variation *σ*_*g*_.
c. Equivalence conclusion of a GM crop (or a GM trait) relies on the totality of evidence across key components when compared a given control background. Codex guidelines state [2] that “A variety of data and information are necessary to assess unintended effects because no individual test can detect all possible unintended effects or identify, with certainty, those relevant to human health. These data and information, when considered in total, provide assurance that the food is unlikely to have an adverse effect on human health”. In practice a comprehensive assessment has been performed over a wide range of endpoints from various studies e.g. often > 50 analytes in a composition study alone, and any experimental deviation from equivalence has been evaluated in terms of the “natural variation” as well as the nominal level of the statistical significance. With such an assessment, any limitation of the evidence due to a given control is minimized and an equivalence of a GM trait if concluded could be assumed for different genotypic background after the commercialization.

In summary, by OECD and WHO/FAO guidelines, GM crop safety assessment is characterized by multiple comparisons of a GM crop across key endpoints with the conventional references with a given (control) genotypic background. Three features of the assessment, focus on the GM trait effect, a wide range of background variations, and multiplicity of the comparisons, are considered in the following section for the derivation of a set of conditional equivalence criteria.

### A conditional equivalence criterion for similarity

Let (*D*_*TC*_, *D*_*CR*_, *D*_*TR*_) denote estimates of (Δ_*TC*_, Δ_*CR*_, Δ_*TR*_) with variances 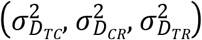., At first, equivalence criteria of EFSA [6] and Vahl and Kang [8] from the current literature were formulated in the notation of this manuscript. Then a conditional equivalence criterion was derived with a mixed model approach and relationships among three types of criteria were discussed.

EFSA equivalence criterion (or limit) was defined by the following equation under EFSA model 2, an ad hoc model assuming both test and control in a TCR trial as random and independent varieties with the same variance as a reference but unspecified means.

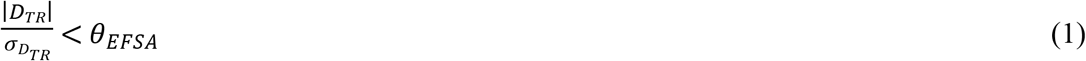

where *D*_*TR*_ as stated repeatedly is a mixture of the GM trait effect and the control background effect, 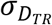 consists of both genetic and residual variations (due to sampling), and *θ*_*EFSA*_ was specified by a 95% confidence limit of a t distribution with a sample estimate of 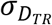 in practice. Clearly, EFSA equivalence criterion considered only the background variation of the test variety (shared with the control) and no GM trait effect which, if presented, would have adopted a non-central t distribution for *θ*_*EFSA*_.

Vahl and Kang’s scaled average equivalence criterion is defined in a similar way but in the scale of the genetic portion of the background variation [8].

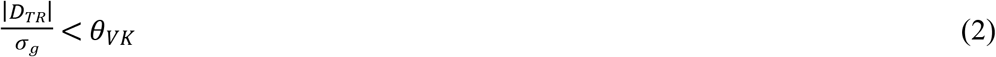

where 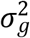 is the leading term of the genetic variation of 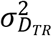 in (1), and *θ*_*VK*_ was specified by a 95% confidence limit of a standard normal. Nevertheless, underlying assumptions, i.e. the GM trait effect is zero and the background of a GM test variety being the same as a random reference, are the same for both criteria (1) and (2).

Let *E*_*E*_ and *E*_*C*_ denote respective expectation with respect to the environmental effect, such as site, replicate and residual, and the control background effect following the distribution of references. A mixed model approach for a fixed effect Δ_*TC*_ and a random effect Δ_*CR*_ assumes *E*_*C*_ [*E*_*E*_(*D*_*TR*_)] = *E*_*C*_[Δ_*TR*_] = *E*_*C*_[Δ_*TC*_ + Δ_*CR*_] = Δ_*TC*_, and 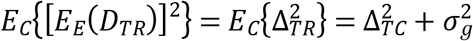. When applying to Vahl and Kang’s scaled average equivalence, the following equation can be derived

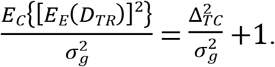

While the first expectation *E*_*E*_(∙) indicates the test statistic and the parameter of interest the same as for criteria (1) and (2), the second expectation *E*_*C*_(∙) reveals the change of hypothesis if a control background were assumed as random. Clearly, Δ_*TR*_ is for an equivalence with the given control background as one part of the assessment, and Δ_*TC*_ is for a marginal equivalence or a conditional equivalence for a random background if absence of an interaction. Since *E*_*E*_ [*E*_*C*_(*D*_*TR*_)] = *E*_*C*_[*E*_*E*_(*D*_*TR*_)] = Δ_*TC*_, *E*_*C*_(*D*_*TR*_) would represent an estimate of Δ_*TC*_, and an equivalence criterion for a GM trait effect could be defined implicitly by

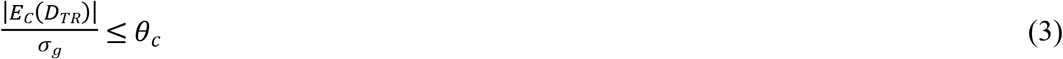

where *θ*_*c*_ is a conditional equivalence criterion to be discussed in the following. This mixed model approach has been applied by FDA for individual equivalence [16] and by Vahl and Kang for a distribution-wise equivalence in GM crop assessment [8]. Therefore, a conditional equivalence is defined solely for a GM trait effect Δ_*TC*_, and a criterion *θ*_*c*_ could be derived as a function of the background variation (Fig 1).

Let *E*_*C*_(*D*_*TR*_) = (*D*_*TR*_│*D*_*CR*_ = 0) denote a conditional difference with a mean 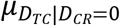 which has been shown to be the best estimator of the trait effect based on the correlation structure among three genotypic group means in a TCR trial [15]. Note that Δ_*CR*_ = 0 is a key assumption in defining the natural variation for GM crop safety assessment. Thus, 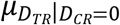 and Δ_*TC*_ are interchangeable in terms of the parameter value (or the equivalence criterion), but a statistic for 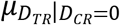 would be applied for an equivalence testing.

Parallel comparisons of criteria (1), (2), and (3) could be made using mean-squares of expected differences i.e. [*E*_*E*_(∙)]^2^ to demonstrate differences in the statistical hypotheses.

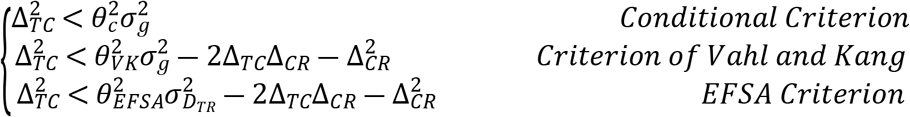

First, by criteria of EFSA and Vahl and Kang, equivalence of a trait effect Δ_*TC*_ would be largely determined by the background effect Δ_*CR*_, not only the magnitude but also its direction. Opposite signs of Δ_*TC*_ and Δ_*CR*_ would be much more likely to be concluded as equivalent than those with the same sign do. In addition, the probability thresholds for *θ*_*EFSA*_ and *θ*_*VK*_ assume Δ_*TC*_ = 0, not a requirement of the principle of substantial equivalence. In contrast, the conditional equivalence criterion does not assume Δ_*TC*_ = 0, but Δ_*TC*_ = 0 is expected to provide a maximum chance of concluding equivalence regardless of the sign and magnitude of Δ_*CR*_ for a given control.

Second, a conditional equivalence standard could be derived as 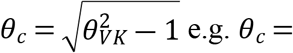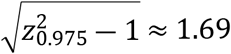 of which an interpretation could be provided as follows. For a GM test with a trait effect |Δ_*TC*_| < 1.69*σ*_*g*_, when averaging over the background effect, the GM test crop would be within the 95% confidence interval of a reference. Therefore, the conditional criterion *θ*_*c*_*σ*_*g*_ is for a trait effect and defined by the range of reference variation thus follows the OECD guidelines [1,2,3].

Third, EFSA equivalence criterion is a much loosely defined criterion as a function of the experimental design (i.e. numbers replicates and sites and total number of references) with *θ*_*EFSA*_ > *θ*_*VK*_ and 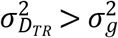, which compromises the efficacy of detecting an unintended effect if presents as pointed out by Vahl and Kang [8, p23], and tends to encourage a trial with lower number of reference and consequently lead to an arbitrary conclusion of equivalence as concerned by Ward et al. [10].

In summary, a conditional equivalence criterion independent of the background was derived in this section in the scale of the reference variation, thus called a variation-scaled criterion. However, while criteria of EFSA and Vahl and Kang are all variation-scaled criteria, if applied as a one-fits-all criterion, certain problems are inevitable. Firstly, even though EFSA classifies those cases with zero estimate of *σ*_*g*_ as “Equivalence Not Concluded”, an arbitrary conclusion is expected as *σ*_*g*_ becomes less than certain threshold (relative to the residual variation) due to a large proportion of close to zero criterion. A second problem is that, with a criterion defined by a 95% confidence limit, false negative (i.e. non-equivalence by chance) is expected to be at least 5% for each endpoint and would be much higher due to the proof-of-equivalence. While a comprehensive assessment requires a totality of evidence, optional criteria become necessary.

## Alternative criteria in a comprehensive assessment

### An empirical mean-scaled criterion when *σ*_*g*_ is low

Alternative criteria are discussed in this section when the reference variation is too low for a variation-scaled criterion. When 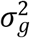 is low, references in a TCR trial become similar to each other including the control. Consequently, the equivalence of a GM crop may become the same as the bioequivalence of a generic drug to a brand-named reference in pharmaceutical industry in terms of the comparison between the test and the control and the absence of references.

The 80-125% rule has long been adopted in pharmaceutical industry for two drugs being “similar to such a degree that their effects, with respect to both efficacy and safety, will essentially be the same” [20,21]. This standard is also recommended for GM crop equivalence assessment in this research. That is, for endpoints with low *σ*_*g*_, a conditional equivalence would be defined in a mean-scale by

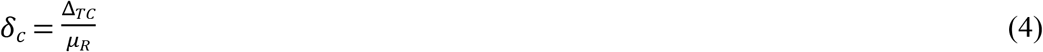

Under the 80-125% rule, standards were differentiated between Δ_*TC*_ < 0 or Δ_*TC*_ > 0. For simplicity *δ*_*c*_ = 0.25 is applied in this research, and under a log-transformation as recommended by FDA, the criterion becomes Δ_*TC*_ = ± log (1 + *δ*_*c*_) =± log (1.25).

While the variation-scaled standard *θ*_*c*_ = 1.69 is based on the concept of equivalence under natural variation with a history-of-safe-use, the mean-scaled standard *δ*_*c*_ = 0.25 is empirical. Two standards appear to be independent in theory, but in practice they are highly correlated as will be shown in the following maize grain composition example. Let *CV*_*g*_ = *σg/μR* denote a coefficient of genetic variation among references in the original scale of a TCR trial data. Two equivalence standards would be equal, i.e. 1.69*σ*_*g*_ = 0.25*μ*_*R*_, at *CV*_*g*_ = 14.8%. When a log-transformation is applied, a log-normal distribution of *y* = *log* (*x*) is assumed and two standards become the same in a log-scale, i.e. 1.69*σ*_*gy*_ = log (1.25), at *CV*_*g*_ = 13.3% using 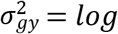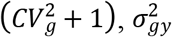 as the reference variance in the log-scale. As will be shown in the example, *CV*_*g*_ = 13.3~14.8% are near the center of the maize grain composition data, and two standards 1.69 *σ*_*g*_ and 0.25*μ*_*R*_ are in fact largely overlapped due to a narrow range of *CV*_*g*_ and a wide separation of means across analytes.

### Optional criteria for multiple comparisons

Equivalence criteria of EFSA and Vahl and Kang in the previous section did not apply a whole range of reference variation, and with a 95% confidence limit a minimum 5% false negative is expected even with no trait effect and could be much higher due to the proof-of-equivalence (as shown in the following example). The same is true for a conditional equivalence. Therefore, an optional criterion 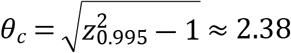 corresponding to a 99% confidence limit is recommended to control the number of false negative.

For a use of whole range of “natural variation” in a proof-of-equivalence, OECD provided summary of some historic data including mean and range of maize and soybean compositions [22,23]. In the meantime, EFSA adopted an intuitive evaluation using box plots of the test, the control and references for analytes failed to conclude equivalence by the equivalence testing.

However, these approaches were unable to separate the true (genetic) “unintended effect” of a GM trait from those due to environments. In contrast, *θ*_*c*_ = 2.38 considers only genetic variation and provides a statistically interpretable standard of equivalence.

In addition, *δ*_*c*_ = 0.5 is also recommended as an optional criterion for endpoints with low reference variation following the practice of pharmaceutical industry for high variant endpoints [24].

### A data-driven procedure for criterion selection

The following diagram describes the proof-of-equivalence approach as requested by EFSA, similar to those in pharmaceutical industry [16,24,25]. The diagram can be applied on a mean of a GM test or a difference between a GM test and references.

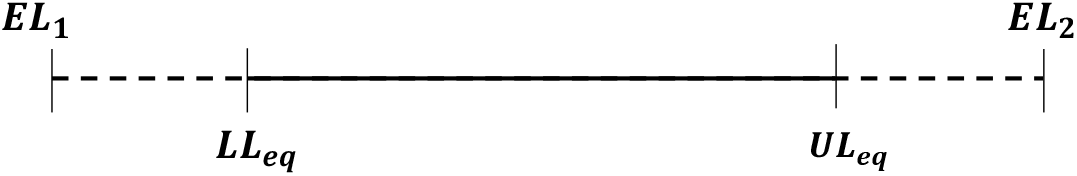

where *EL*_1_ and *EL*_2_ are lower and upper limits of an equivalence criterion, (*LL*_*eq*_, *UL*_*eq*_) is a (confidence) interval containing critical values of the estimated mean or difference for concluding equivalence. The dash lines consist of the estimated mean or difference failed to be concluded as equivalent, called the burden of proof-of-equivalence, due to the margin-of-error in comparisons with *EL*_1_ and *EL*_2_ at the designated significance level.

Assume a TCR trial with eight sites each of four replicates per site/treatment currently requested by EFSA and generally accepted by most international regulatory agencies. The trait effect is best estimated by a conditional difference with a mean 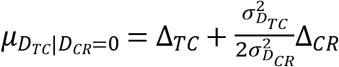 and a variance 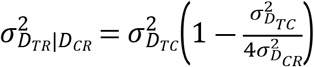 as derived in Jiang et al. [15]. For simplicity, no genotype by environmental interactions were assumed. With a given *σ*_*g*_:*σ*_*e*_ (or a given *σ*_*g*_ at *σ*_*e*_ = 1), a normal approximation was applied in the following for an asymptotic equivalence analysis using an interval (*LL*_*eq*_, *UL*_*eq*_) as functions of (*EL*_1_*, EL*_2_) defined by equations (3) and (4).

Let *EL*_*c*_ = *EL*_2_ = ―*EL*_1_ for a GM trait effect. In this section, a threshold of *σ*_*g*_:*σ*_*e*_ for alternating *EL*_*c*_ = 1.69*σ*_*g*_ and 0.25*μ*_*R*_, a key question in criterion selection, is investigated. Obviously, no equivalence could be concluded even for Δ_*TC*_ = 0 with *EL*_*c*_ = 1.69*σ*_*g*_ if *σ*_*g*_:*σ*_*e*_ is close to zero, and the threshold of *σ*_*g*_:*σ*_*e*_ should be large enough for *EL*_*c*_ = 1.69*σ*_*g*_ to provide an 80% power of equivalence under a regulatory requested design. Fig 2 presents numerical results of an asymptotic equivalence analysis (ignored the variation in estimating *σ*_*g*_). Variation settings *σ*_*g*_:*σ*_*e*_ = (0 ~ 3) in the left plot and the residual coefficient of variation *CV*_*e*_ = (0 ~ 30%) in the right plot is based on the maize grain composition example in the following section. In the left plot, a grid search was performed over Δ_*TC*_ = (0 ~ 2.38*σ*_*g*_) for each value of *σ*_*g*_:*σ*_*e*_, the maximum trait effect for 80% power of equivalence was obtained with *EL*_*c*_ = 1.69*σ*_*g*_ and 2.38*σ*_*g*_ as functions of *σ*_*g*_:*σ*_*e*_. Similarly, in the right plot is for *EL*_*c*_ = 0.25*μ*_*R*_ and 0.5*μ*_*R*_ as functions of *CV*_*e*_.

**Fig 2.**
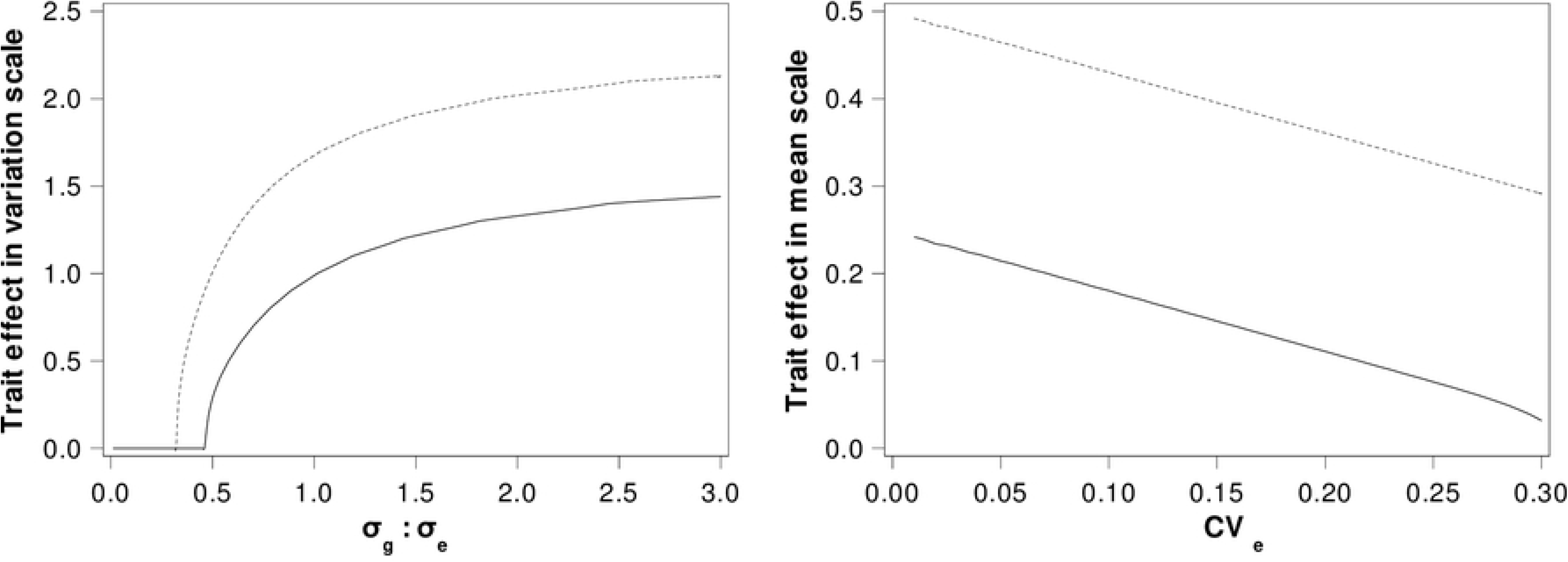
Asymptotic maximum GM trait effects for 80% power of equivalence under EFSA requested design. Left plot: Asymptotic maximum GM trait effects with *EL*_*c*_ = 1.69*σ*_*g*_ and 2.38 *σ*_*g*_ as functions of *σ_g_*:*σ_e_*; Right plot: Asymptotic maximum GM trait effects with *EL_c_* = 0.25_*μR*_ and 0.5*μ*_*R*_ as functions of residual coefficient of variation *CV*_*e*_.

Results in Fig 2 indicate that, in general, an 80% power of equivalence requires a trait effect Δ_*TC*_ substantially less than *EL*_*c*_ due to the proof-of-equivalence. Asymptotically, a threshold *σ*_*g*_: *σ*_*e*_ = 1.0 could be applied for alternating *EL*_*c*_ = 1.69*σ*_*g*_ and 0.25*μ*_*R*_ (Fig 2). When *σ*_*g*_:*σ*_*e*_ ≥ 1.0, the maximum trait effect for equivalence is about 1.4*σ*_*g*_ for *EL*_*c*_ = 1.69*σ*_*g*_ and 2.1*σ*_*g*_ for *EL*_*c*_ = 2.38*σ*_*g*_. When *σ*_*g*_:*σ*_*e*_ < 1.0, even a negligible trait effect might not have enough power to conclude equivalence by either criterion. By *EL*_*c*_ = 0.25*μ*_*R*_, the maximum trait effect for equivalence is about 0.15*μ*_*R*_ when *CV*_*e*_ = 15%. Note that Fig 2 did not include the variation in estimating *σ*_*g*_ which should be a function of the number of references and, when considered, the estimated maximum trait effect in Fig 2 could be substantially lower.

Results in Fig 2 demonstrate no existence of a one-fits-all criterion under practical ranges of *σ*_*g*_:*σ*_*e*_ and *CV*_*e*_. Therefore, the following set of conditional equivalence criteria was proposed with a three-step procedure for criterion selection.

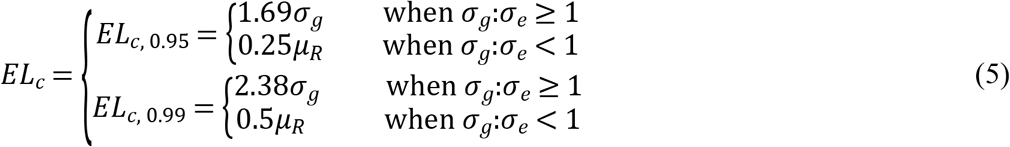

Firstly, *EL*_*c*,0.95_ = 1.69*σ*_*g*_ is applied as the primary criterion whenever a reasonable variation *σ*_*g*_ is available, i.e. *σ*_*g*_:*σ*_*e*_ ≥ 1. Secondly, an alternative *EL*_*c*,0.95_ = 0.25*μ*_*R*_ is used for *σ*_*g*_:*σ*_*e*_ < 1. Thirdly, *EL*_*c*, 0.99_ is optional for endpoints failed to show equivalence with *EL*_*c*, 0.95_ expectedly only for a small proportion of endpoints say 5 to 10% or less.

In practice, the procedure (5) depends on estimates of *σ*_*g*_, *μ*_*R*_, and *σ*_*g*_:*σ*_*e*_ and variations of these estimates will be a function of the experimental design.

## Application to a maize grain composition example

Maize grain composition data in Jiang et al. [15], originally applied by EFSA for method demonstration, were reanalyzed with and without the log transformation. At first, reference means and variations were estimated from 13 references for each analyte, and paired estimates across all 53 analytes for *EL*_*c*_ = (1.69*σ*_*g*_, 0.25*μ*_*R*_) without transformation and *EL*_*c*_ = (1.69*σ*_*g*_, log (1.25)) with transformation were plotted (Fig 3). In the left plot with no transformation, a strong linear correlation can be observed between estimates of *σ*_*g*_ and *μ*_*R*_ (with an estimated *r*^2^ = 0.9644 in the log-scale). The observed *CV*_*g*_ (= *σ*_*g*_/*μ*_*R*_) is highly consistent within a range (0 ~ 25.2%) and a mean 10.2% across a wide range of *μ*_*R*_. The observed *CV*_*e*_ has a mean 7.2% and a range (0.64 ~ 28.2%). In the right plot with the log transformation, *EL*_*c*_ = 1.69*σ*_*g*_ indeed tends to be independent of the mean. The mean estimate of *EL*_*c*_ = 1.69*σ*_*g*_ is 0.17, slightly lower than the line *EL*_*c*_ = *log*(1.25) ≈ 0.22 in the plot.

**Fig 3.**
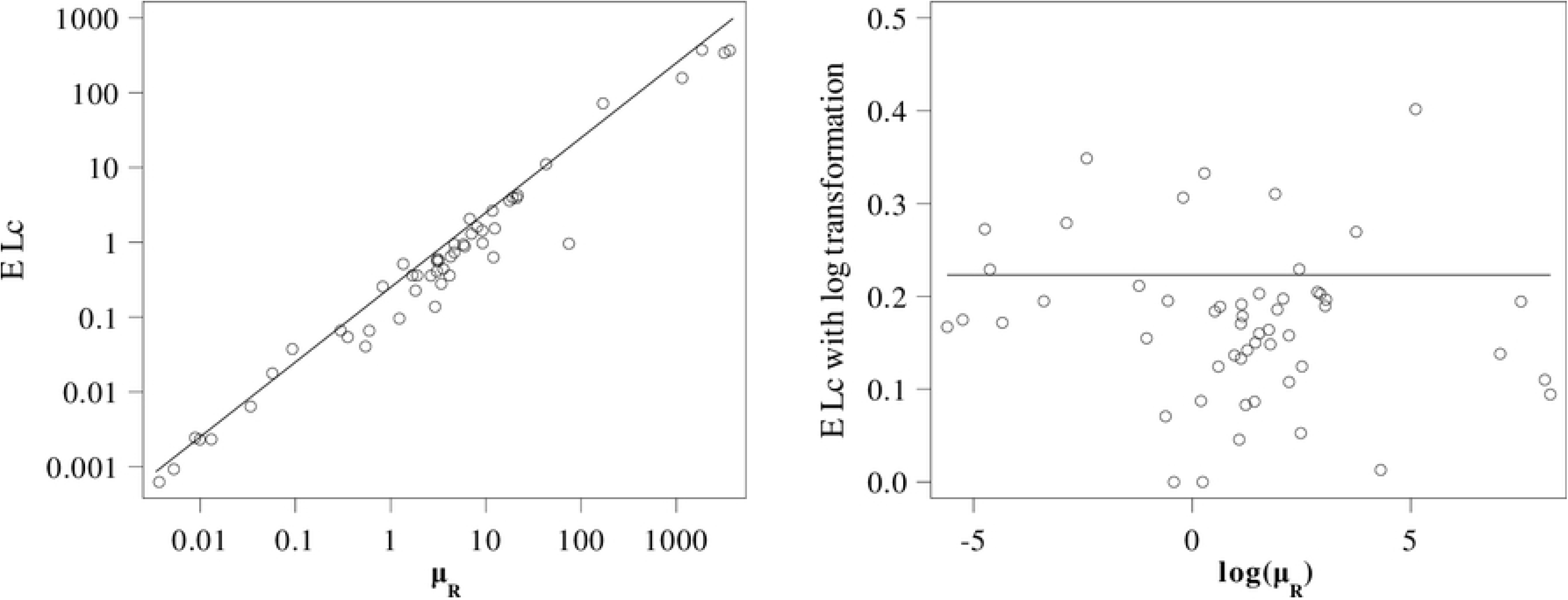
EFSA example: Estimated equivalence criteria *EL*_*c*_ = 1.69*σ*_*g*_ (markers) and 0.25*μ*_*R*_ (line) across 53 analytes. Left plot: Without transformation; Right plot: With log transformation.

Results from Fig 3 have at least two implications. Firstly, high comparability between equivalence criteria *EL*_*c*_ = (1.69*σ*_*g*_, 0.25*μ*_*R*_) strongly supports the equivalence evaluation of a GM trait effect as a similarity measurement for a whole range of the reference variation. While the concept of substantial equivalence is generally understood as comparing a GM crop with the reference variation, a mean-scaled criterion in the original unit is a natural alternative when *σ*_*g*_ is small and lack of a good estimate. Secondly, *EL*_*c*_ = 1.69*σ*_*g*_ on average appears to be more conservative than *EL*_*c*_ = 0.25*μ_R_*, an empirical support for *EL*_*c*_ = 1.69*σ*_*g*_ in terms of mean percent difference.

Table 1 summarizes the estimate of 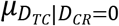 in scales of *σ*_*g*_ and 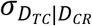 (i.e. the same scale as the criterion) and *σ*_*D*_*TC*|*D_CR_* (i.e. the t value of a conditional difference test by results of EFSA model 1 and model 2). Values in bold follow the procedure (5). Though only two analytes with zero estimate for *σ*_*g*_, 9 and 10 analytes are estimated as *σ*_*g*_:*σ*_*e*_ < 1 when no transformation or transformation was applied.

**Table 1.** Summary of estimates of 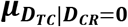 in ratios to *σ*_*g*_, *μ*_*R*_, and 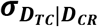 (results following the procedure (5) marked as bold) and comparisons of three criteria with difference *D* = *D*_*TR*_ for EFSA and Vahl and Kang (VK) methods, and 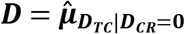 for conditional equivalence (CD).

| $\sigma_g:\sigma_e$ | Mean estimate and range of $\vert \mu_{D_{TC}\vert D_{CR}=0}\vert $ in ratios to | | | | # Analytes<br>(with $\vert D\vert > EL$) | | |
| --- | --- | --- | --- | --- | --- | --- | --- |
| | #<br>Analyte | $\sigma_g$ | $\mu_R$ | $\sigma_{D_{TC}\vert D_{CR}}$ | EFSA | VK | CD |
| In original unit |  |  |  |  |  |  |  |
| $\geq 1$ | 44 | **<u>0.34(0.00~1.12)</u>** | 0.04(0.00~0.13) | 1.48(0.01~5.25) | 2 | 3 | 0 |
| $< 1$ | 9 | 1.35(0.59~2.12)* | **<u>0.09(0.02~0.19)</u>** | 1.90(0.93~3.80) | 1* | 2* | 0 |
| With log transformation |  |  |  |  |  |  |  |
| $\geq 1$ | 43 | **<u>0.37(0.01~1.24)</u>** | 0.04(0.00~0.14) | 1.61(0.03~6.47) | 2 | 5 | 0 |
| $< 1$ | 10 | 1.02(0.17~1.81)* | **<u>0.09(0.02~0.22)</u>** | 1.69(0.35~3.79) | 1* | 1* | 0 |

\*: Two analytes with zero estimate of $\sigma_g$ are not included.

While large t values for estimates of 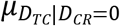 in the ratio to 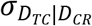 are evidence of non-zero trait differences, results in bold demonstrate the good performance of the procedure (5). For example, for those analytes with *σ*_*g*_:*σ*_*e*_ < 1 and *EL*_*c*_ = 0.25*μ*_*R*_ without transformation or *EL*_*c*_ = log (1.25) with transformation, estimates of 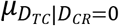 are all well within the limit (i.e. 0.25) in the scale of *μ*_*R*_, but several of them would have exceeded the limit (i.e. 1.69) if *EL*_*c*_ = 1.69*σ*_*g*_ was applied. Results are highly consistent with and without transformation.

The last three columns of Table 1 summarize comparisons of the estimated difference with three criteria: EFSA, VK for Vahl and Kang, and CD for the conditional equivalence. With a proof-of-equivalence, an estimated difference exceeding *EL* would automatically lead to a non-equivalence conclusion. |*D*_*TR*_| > *EL* were observed for both criteria of EFSA and VK. Three cases of |*D*_*TR*_| > *EL* for EFSA criterion represents almost exactly a 5% of non-equivalence by chance, 2.6 (i.e. 5% of 51) in expectation which simply suggests no evidence of GM trait effects. These results are the same as EFSA original analysis with the transformation, where three analytes were classified as “Non-Equivalence More Likely Than Not” or “Non-Equivalence”. Yet a total of nine analytes (i.e. 17% of 53) failed to conclude equivalence including two with zero estimate of *σ*_*g*_, much higher than the nominal 5% level due to the proof-of-equivalence. However, for the conditional criteria no 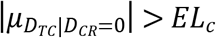 was observed. Note that although no trait effect exceeds *EL*_*c*,0.95_ in Table 1, *EL*_*c*,0.99_ might still be necessary in a formal testing due to the proof-of-equivalence.

In summary, despite of a formal statistical testing yet to be developed, simple comparisons in the example demonstrate obvious advantages of the conditional equivalence criteria proposed in this investigation over those of EFSA and Vahl and Kang in a comprehensive equivalence assessment.

## Discussion

Under OECD guidelines GM crop safety assessment is characterized by the comparative approach on a GM crop of known (control) genotypic background to conventional references with a history-of-safe-use. The equivalence testing method prescribed by EFSA assesses differences between the GM variety and a group of commercial reference varieties. These differences, as discussed by Jiang et al. [9], may be driven by a trait effect, a known control background effect, or both. The EFSA equivalence criterion consists of only the background variation, and three direct consequences are worth noting. First, the EFSA criterion is entirely for the random background effect, contradicting with the principle focusing on “unintended effect” of a GM trait. Second, the EFSA criterion becomes degenerate when the reference variation is low and estimated as zero or close to zero, a common case in composition studies. Third, it is the inability to control the false negative rate, i.e. substantially higher than the target 5% level as defined by criteria of EFSA [6] or Vahl and Kang [8] due to the proof-of-equivalence even in the absence of a true GM trait effect.

A set of conditional equivalence criteria under the same assumptions as those of EFSA and Vahl and Kang are derived in this manuscript. However, the new criteria are for a GM trait effect Δ_*TC*_, which is independent of the genotypic background of a given control and thus fundamentally different from those of EFSA and others.

The approach using the reference variation *σ*_*g*_ as a scale in this manuscript, if available, follows the principle of substantial equivalence of OECD guideline that “any observed differences should be assessed in the context of the range of natural variations” [2]. When the reference variation *σ*_*g*_ is small, an alternative criterion was proposed to apply the scale of the reference mean *μ*_*R*_ with the procedure (5). The parallel nature of the mean-scaled and the variation-scaled criteria lies in the definition of equivalence to a fixed reference (with a mean percentage difference as an empirical standard) or to a group of references (with a fold of standard deviation defining the range of a history-of-safe-use). Re-analysis of the maize grain composition example originally applied by EFSA illustrate that even though only two endpoints have *σ*_*g*_ estimated as zero, low values of *σ*_*g*_ relative to the residual are common (about 20% in the example by the threshold *σ*_*g*_:*σ*_*e*_ < 1). In these cases, a mean-scaled criterion as defined in (5) should be considered as a natural alternative to a variation-scaled criterion, which is strongly supported by the close correlation between the mean and variation in the example (Fig 3) and commonly observed in biological literature (e.g. the log-transformation suggested by EFSA and FDA in pharmaceutical studies).

Another type of criterion in the literature for endpoints with low values of *σ*_*g*_:*σ*_*e*_ could be labeled as “phenotypic equivalence” due to including residual variation *σ*_*e*_ (and other environmental variations) as part of the “natural variation” in defining equivalence. One example is the distribution-wise equivalence proposed by Vahl and Kang [8] and applied by Van der Voet et al. [26] in the analysis of five studies in which GM crops were fed to rats. Another example is the criterion applied by Schmidt et al. [27], based on one unit of reference standard deviation, i.e.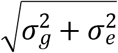 in expectation, following an example in the EFSA guideline [28]. An intuitive interpretation of the “phenotypic equivalence” would be a large proportion of overlapping between observed responses of the test and the control. However, no discussion could be found on the level of “unintended effect” of a GM trait in the unit of *σ*_*e*_, either in terms of the regulatory policy or a biological interpretation. In addition, practical implications of these criteria would depend on the type of the study (e.g. the applicable sample size) and the characteristic of the endpoint (e.g. magnitudes of *CV*_*g*_ and *CV*_*e*_).

In their guideline, EFSA also proposed a simulation approach for evaluating equivalence by an empirical distribution of the number of significant outcomes in the difference testing between two independent references. Regardless of the residual variation, the absolute mean difference between two references is 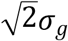 and a 95% confidence limit would be approximately 2.8*σ*_*g*_ under normality. From this perspective, *EL*_*c*_ = 1.69*σ*_*g*_ and 2.38*σ*_*g*_ in (5) would be considered as conservative. Therefore, even though, under certain circumstances, some assumptions would be more plausible than others, the comparative assessment of GM crops should be a comprehensive approach with false negatives under control.

A false negative in an equivalence testing is in many ways similar to the false positive in a difference test. In the maize grain composition example, the EFSA criterion demonstrated an almost exact 5% of the analytes with observed differences greater than the equivalence limit (i.e. 3/51 or 5.9%) (Table 1). However, due to the proof-of-equivalence approach, a much higher proportion of analytes (i.e. 9/53 or 17%) failed to conclude equivalence by the EFSA method [6]. However, all analytes in the example are within the conditional limits *EL*_*c*,0.95_ = 1.69*σ*_*g*_ or 0.25*μ*_*R*_. In addition, a step-wise procedure with optional criteria *EL*_*c*,0.99_ = 2.38*σ*_*g*_ or 0.5*μ*_*R*_ was proposed in this investigation for endpoints failed to conclude equivalence at *EL*_*c*,0.95_. In contrast to use the box plots of EFSA or historic data [22,23,29], criteria *EL*_*c*,0.99_ consist of no environmental effects, thus are true criteria of “unintended effect” due to multiple comparisons. References in the current TCR trial though limited in number often may be a more reliable source of information due to difficulties in estimating the genotypic variation among references from historic data.

With the criteria developed here, an immediate further research subject would be statistical methods of conditional equivalence testing applying these criteria. Because these criteria must be estimated from the reference data and the variation of the estimation must be taken into account especially when the number of references is limited. The variation of the estimated criterion would be compounded with those of the trait effect, not accounted for in the EFSA equivalence testing, which partially contributed to its poor performance.

## Acknowledgments

We thank Duška Stojsin and John Vicini for providing constructive feedback on the manuscript.

## Supporting information

**S1 Fig. Graphical illustration of EFSA and conditional equivalence criteria.** EFSA equivalence criterion is defined primarily by a (random) control background effect and has no specification of a GM trait effect, and a conditional equivalence criterion is defined solely for a GM trait effect with a given control.

**S2 Fig. Asymptotic maximum GM trait effects for 80% power of equivalence under EFSA requested design.** Left plot: Maximum GM trait effects with *EL*_*c*_ = 1.69*σ*_*g*_ and 2.38*σ*_*g*_ as functions of *σ*_*g*_:*σ*_*e*_; Right plot: Maximum GM trait effects with *EL*_*c*_ = 0.25*μ*_*R*_ and 0.5*μ*_*R*_ as functions of residual coefficient of variation *CV*_*e*_.

**S3 Fig. EFSA example: Estimated equivalence criteria of** *EL*_*c*_ = 1.69*σ*_*g*_ **(markers) and** 0.25*μ*_*R*_ **(line) across 53 analytes.** Left plot: Without transformation; Right plot: With log transformation.

## References

1. OECD, Safety evaluation of foods derived by modern biotechnology: Concepts and principles. 1993, Organisation for economic co-operation and development: Paris, France.

2. Codex Alimentarius, Guideline for the conduct of food safety assessment of foods derived from recombinant-DNA plants. 2003, Codex Alimentarius Commission, Joint FAO/WHO Food Standards Programme, Food and Agriculture Organization of the United Nations: Rome, Italy.

3. Codex Alimentarius, Foods derived from modern biotechnology. 2009, Codex Alimentarius Commission, Joint FAO/WHO Food Standards Programme, Food and Agriculture Organization of the United Nations: Rome, Italy.

4. FAO. GM Food Safety Assessment: Tools for traners. Food and Agriculture Organization of the United Nations Rome, 2008. ISBN 978-92-5-105978-4.

5. EFSA, Commission Implementing Regulation (EU) No 503/2013 of 3 April 2013 on. for authorisation of genetically modified food and feed in accordance with Regulation (EC) No 1829/2003 of the European Parliament and of the Council and amending Commission Regulations (EC) No 641/2004 and (EC) No 1981/2006 Text with EEA relevance. 2013, European Food Safety Authority: Parma, Italy.

6. EFSA, Scientific opinion on statistical considerations for the safety evaluation of GMOs. EFSA Journal, 2010. 8: p. 1–59.

7. Kang Q and Vahl CI. Statistical analysis in the safety evaluation of genetically-modified crops: Equivalence tests. Crop Science, 2014. 54: p. 2183–2200.

8. Vahl CI and Kang Q. Equivalence criteria for the safety evaluation of a genetically modified crop: A statistical perspective. The Journal of Agricultural Science, 2015. 154(03): p. 383–406.

9. Jiang C, Meng C, Schapaugh A. Comparative analysis of genetically-modified crops: Part 1. Conditional difference testing with a given genetic background. PLoS ONE, 2019. 14(1): e0210747. https://doi.org/10.1371/journal.pone.0210747

10. van der Voet H, Perry JN, Amzal B, and Paoletti C. A statistical assessment of differences and equivalences between genetically modified and reference plant varieties. BMC Biotechnology, 2011. 11(15): p. 1–20.

11. Ward KJ, Nemeth MA, Brownie C, Hong B, Herman RA, and Oberdoerfer R. Comments on the paper “A statistical assessment of difference and equivalences between genetically modified and reference plant varieties” by van der Voet et al. 2011. BMC Biotechnology, 2012. 12(13): p. 1–7.

12. Fernandez A and Paoletti C. Unintended Effects in Genetically Modified Food/Feed Safety: A Way Forward. Trends in Biotechnology, 2018. 36(9): p. 872–875.

13. Simó C, Ibáñez C, Valdés A, Cifuentes A. and García-Cañas V. Metabolomics of Genetically Modified Crops. Int. J. Mol. Sci. 2014. 15: p. 18941–18966; Doi:10.3390/ijms151018941

14. Herman RA, Fast BJ, Scherer PN, Brune AM, de Cerqueira DT, Schafer BW, et al. Stacking transgenic event DAS-O15O7-1 alters Herman RA less than traditional breeding. Plant Biotechnology Journal, 2017. 15(10): p. 1264–1272.

15. Herman RA, Huang E, Fast B, and Walker C. EFSA Genetically Engineered Crop Composition Equivalence Approach: Performance and Consistency. Journal of Agricultral and Food Chemistry, 2019. 67(14): p. 4080–4088.

16. FDA, Guidance for industry: Statistical approaches to establishing bioequivalence. 2001, U.S. Department of Health and Human Services, Food and Drug Administration, Center for Drug Evaluation and Research.

17. Prado JR, Segers G, Voelker T, Carson D, Dobert R, Phillips J, et al. Genetically Engineered Crops: From Idea to Product. Annu. Rev. Plant Biol. 2014. 65: p. 769–90.

18. Venkatesh TV, Cook K, Liu B, Perez T, Willse A, Tichich R, et al. Compositional differences between near-isogenic GM and conventional maize hybrids are associated with backcrossing practices in conventional breeding. Plant Biotechnology Journal 2015. 13: p. 200–210.

19. Horak MJ, Rosenbaum EW, Woodrum CL, Martens AB, Mery RF, Cothren JT, et al. Characterization of Roundup Ready Flex Cotton, ‘MON 88913’, for use in ecological risk assessment: Evaluation of seed germination, vegetative and reproductive growth, and ecological interactions. Crop Sci 2007. 47: p. 268–77; http://dx.doi.org/10.2135/cropsci2006.02.0063

20. Schall R and Endrenyi L. Bioequivalence: tried and tested. Cardiovasc J Afr 2010. 21(2): p. 69–70.

21. Kumar SA, Sanjita D. A Review article on Bioavailability and Bioequivalence Studies. International Journal of PharmTech Research 2013. 5: p. 1711–1721.

22. OECD Environmental Health and Safety Publication Series on the Safety of Novel Foods and Feeds. No. 6 Consensus Document on Compositional Considerations for New Varieties of Maize (*Zea Mays*): Key Food and Feed Nutrients, Anti-nutrients and Secondary Plant Metabolites. 2002. OECD Environment Directorate, Paris.

23. OECD Environmental Health and Safety Publication Series on the Safety of Novel Foods and Feeds. No. 25 Revised Consensus Document on Compositional Considerations for New Varieties of SOYBEAN [*Glycine max* (L.) Merr]: Key Food and Feed Nutrients, Anti-nutrients, Toxicants, and Allergens. 2012. OECD Environment Directorate, Paris.

24. Davit BM, Chen ML, Correr DP, Haidar SH, Kim S, Lee CH, et al. Implementation of a Reference-Scaled Average Bioequivalence Approach for Highly Variable Genetic Drug Products by the US Food and Drug Administration. The AAPS Journal 2012. 14(4): p. 915–924.

25. Hsu J, Hwang JTG, Liu HK and Ruberg SJ. Confidence intervals associated with tests for bioequivalence. Biometrika 1994. 81: p. 103–114.

26. van der Voet H, Goedhart PW, Schmidt K. Equivalence testing using existing reference data: An example with genetically modified and conventional crops in animal feeding studies. Food and Chemical Toxicology 2017. 109: p. 472–485.

27. Schmidt K, Schmidtke J, Schmidt P, Kohl C, Wilhelm R, Schiemann J, et al. Variability of control data and relevance of observed group differences in five oral toxicity studies with genetically modified maize MON810 in rats. Arch Toxicol 2017. 91: p. 1977–2006.

28. EFSA Guidance on conducting repeated−dose 90-day oral toxicity study in rodents on whole food/feed. EFSA J 2011. 9: p. 2438.

29. Herman RA, Scherer PN, Phillips AM, Storer NP, Krieger M. Safety composition levels of transgenic crops assessed via a clinical medicine model. Biotechnol. J. 2010. 5(2): p. 172–182.

